# TaxisPy: A Python-based Software for the Quantitative Analysis of Bacterial Chemotaxis

**DOI:** 10.1101/2020.02.06.937714

**Authors:** Miguel Á. Valderrama-Gómez, Rebecca A. Schomer, Michael A. Savageau, Rebecca E. Parales

## Abstract

Several species of bacteria are able to modify their swimming behavior in response to chemical attractants or repellents. Methods for the quantitative analysis of bacterial chemotaxis such as quantitative capillary assays are tedious and time-consuming. Computer-based video analysis of swimming bacteria represents a valuable method to directly assess their chemotactic response. Even though multiple studies have used this approach to elucidate various aspects of the bacterial chemotaxis, to date, no computer software for such analyses is freely available. Here, we introduce TaxisPy, a Python-based software for the quantitative analysis of bacterial chemotaxis. The software comes with an intuitive graphical user interface and can be easily accessed through Docker on any operating system. Using a video of freely swimming cells as input, TaxisPy estimates the culture’s average tumbling frequency over time. We demonstrate the utility of the software by assessing the effect of different concentrations of the attractant shikimate on the swimming behavior of *Pseudomonas putida* F1 and by capturing the adaptation process that *Escherichia coli* undergoes after being exposed to L-aspartate.

## 1. Introduction

Environmental conditions such as temperature, light intensity and chemical composition influence the ability of an organism to grow, reproduce and survive. Many bacteria have evolved intricate mechanisms to sense these and other external stimuli. These signals are processed by the cell and directly affect the functioning of its motility apparatus, thus leading to an informed displacement towards a beneficial environment or away from unfavorable conditions. Chemotaxis refers to a modification in the motility pattern of an organism in response to a change in the chemical composition of its environment. Flagellated bacteria such as *Escherichia coli* and *Pseudomonas putida* swim in liquid environments by rotating their helical flagella. A change in the swimming direction is achieved by the intermittent change of the direction of flagellar rotation, which leads to tumbles that usually last only a fraction of a second (Webre et al. 2003). The swimming pattern of a cell can be quantitatively characterized by its tumbling frequency (Alon et al. 1999, Pohl et al. 2017). In the absence of chemical attractants, the swimming pattern of motile bacteria is described as a “random walk”, which is characterized by short periods of smooth swimming (often termed “runs”) and a high tumbling frequency, with typical values of 0.44 changes of direction per second for *E. coli* (Alon et al. 1999, Staropoli and Alon 2000). After a chemical attractant is sensed, chemotactic bacteria decrease their tumbling frequency to values lower than 0.05 per second (Alon et al. 1999). As a result, cells in a culture effectively swim towards higher concentrations of the attractant following straight paths.

Over the last decades, qualitative and quantitative methods for the study of bacterial chemotaxis have been developed. Some representative protocols include capillary assays (Pfeffer 1884, Adler 1969, Adler 1973), chemical-in-plug assays (Tso and Adler 1974) and computer-based video analyses of swimming cells (Berg and Brown 1972). See Ditty and Parales 2015 for an overview of these methods. While qualitative methods are fast and usually do not involve major technical difficulties, quantitative methods such as the quantitative capillary assay are tedious and time-consuming. Computer-based video analysis of swimming bacteria represents a valuable method to directly and quantitatively assess the chemotactic response of bacteria. Even though multiple studies have used this approach to elucidate various aspects of bacterial chemotaxis (Harwood et al. 1989, Alon et al. 1998, Alon et al. 1999, Staropoli and Alon 2000), to date, no computer software for such analyses is freely available. Here, we introduce TaxisPy, a Python-based software for the quantitative analysis of bacterial chemotaxis. The software comes with an intuitive graphical user interface that is especially suited for users with limited programming knowledge. It systematically addresses typical difficulties associated with the customized use of cell tracking software such as filtering of atypical cell trajectories and determination of parameter values for the identification of tumbles. TaxisPy can be easily accessed through Docker on any operating system.

## 2. Materials and Methods

### 2.1 Bacterial strains, media composition and attractants

The responses of *E. coli* K12 strain RP437 [*thr leu his metF eda rpsL thi ara lacY xyl tonA tsx*] (Parkinson 1978) and *Pseudomonas putida* F1 –both motile organisms used as models for studying chemotactic behavior– to chemical attractants were examined in this study. All cells were grown in Luria-Bertani medium (LB; Sambrook et al. 1989) or minimal medium (MSB; Stanier et al. 1996) at 30°C in a shaking incubator.

### 2.2 Behavioral assays

*Pseudomonas putida* cultures were grown for approximately 18 hours in minimal MSB medium containing 10 mM succinate at 30 °C. One hundred microliters of the culture were sub-cultured into 15 mL minimal MSB containing 10 mM succinate and 5 mM shikimate for induction of chemotactic behavior towards aromatic acids (Luu 2015). After approximately 6 hours of growth, the cells were harvested at mid-exponential phase (OD_600_ = 0.4-0.6) by centrifugation at 5,000 rpm for 10 minutes at room temperature. The cells were gently washed in 15 ml of chemotaxis buffer (CB; 50 mM potassium phosphate buffer pH 7.0, 10 µM disodium EDTA, 0.05% glycerol) and re-centrifuged for 10 minutes at 5,000 rpm (Parales et al. 2000). Finally, the cells were resuspended in 15 mL of aerated CB. For analysis of the swimming pattern of *P. putida*, 10 μl of the cell suspension was placed on a glass slide (Fisher Scientific) and mixed with 1 μl of various concentrations of shikimate in CB to attain final concentrations of 0 μM, 10 μM, 50 μM, 100 μM, and 1 mM. Two seconds after the addition of the attractant, video recording was initiated. Recording of the samples was performed with an Infinity Lite microscope camera (Lumenera, Ottawa, ON, Canada) mounted to a Nikon eclipse TE2000-S inverted microscope, using a magnification of 400X. The swimming behavior was recorded for 1 minute at a speed of 20.3 frames per second using the software Infinity Capture 6.5.4. These videos were then analyzed with TaxisPy as described below to estimate the tumbling frequency of the cell populations.

Samples for analyzing the adaptation of *E. coli* after stimulation with L-aspartate were generated using a different set-up. Cells were grown overnight in LB medium before being harvested by centrifugation (10,000 RPM, 1 minute) and washed with an equal volume of MSB medium. Cells were resuspended in an equal volume of MSB medium. Two hundred microliters of washed culture were inoculated into 15 mL of minimal MSB medium containing 12 mM glucose. When the cultures reached mid-exponential phase (OD_600_= 0.4-0.6), 1 mL aliquots of culture were harvested by centrifugation at 5,000 rpm for 10 min at room temperature and gently resuspended in 1 mL of CB. At regular time intervals, 10 μL samples of *E. coli* were removed and observed with a Nikon eclipse TE200-S inverted microscope at 400X magnification. Using the Infinity Lite mounted camera, 10 second videos were recorded at a speed of 20.3 frames per second. The chemotactic response was initiated by adding L-aspartate to the resuspended cultures at final concentration of 1 mM. To keep cells motile during the experiment, cells were incubated at 30°C in a shaking incubator (200 rpm). These videos were then analyzed with TaxisPy as described below to estimate the tumbling frequencies of the cell populations.

### 2.3 Software and General Workflow

TaxisPy integrates several Python packages to enable the estimation of the tumbling frequency of a bacterial culture (Fig. 1). Its intuitive graphical user interface is based on Jupyter widgets and contains six different tabs (Fig. 2). In order to streamline its distribution, especially among users with limited programming knowledge, TaxisPy can be accessed via a Docker image. Docker is a computer program that performs operating-system-level virtualization and is used to run software packages called containers. Containers bundle their own application, tools, libraries and configuration files and are created from images. Docker has been increasingly used to distribute software across different operating systems in a fast and secure way. To download TaxisPy via Docker, four steps are required:

**Figure 1.**
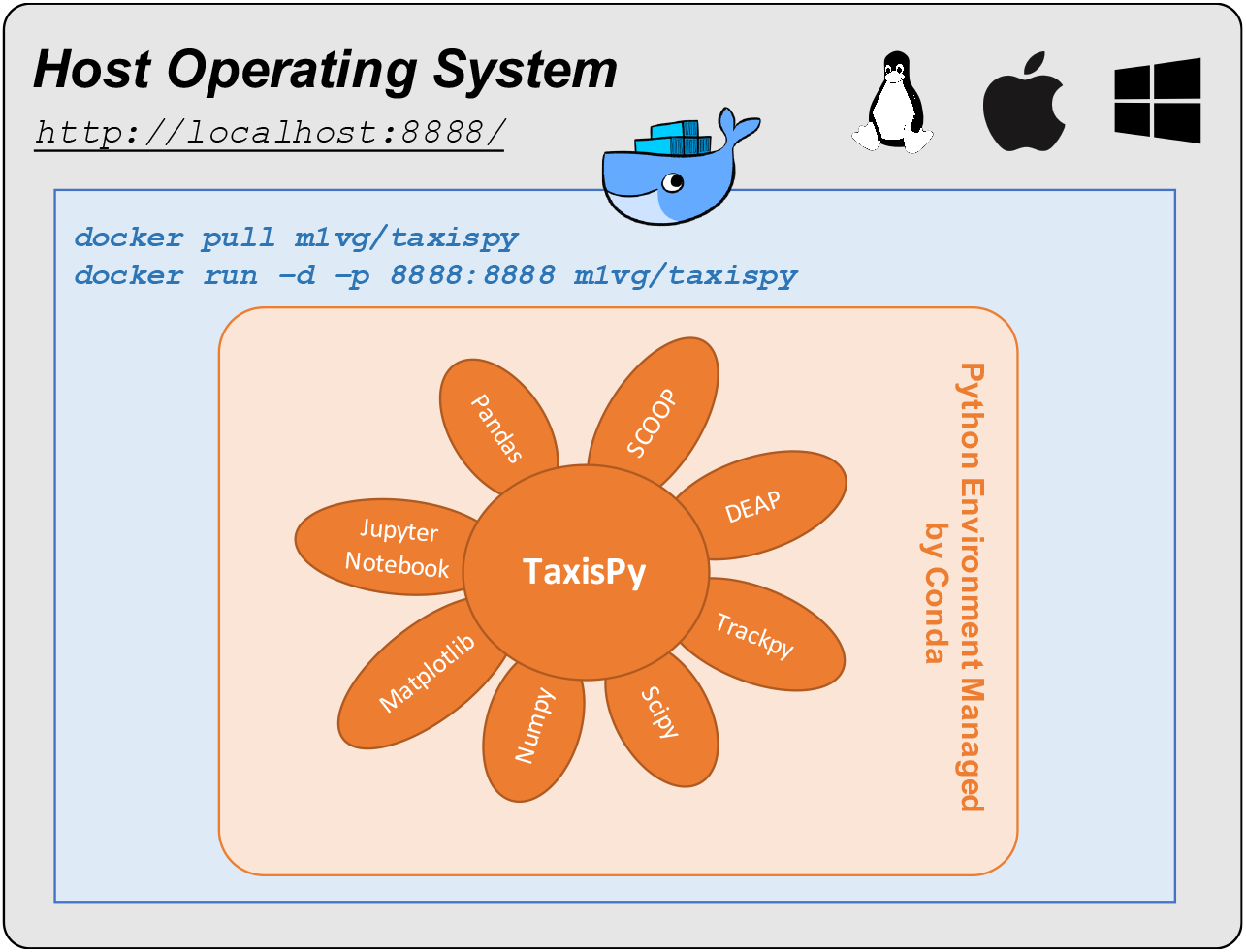
Software architecture. TaxisPy bundles several Python packages that together allow a computer-assisted, quantitative analysis of bacterial chemotaxis. TaxisPy’s user interface is based on Jupyter widgets and can be accessed from Jupyter Notebooks on any web browser, e.g., Google Chrome. TaxisPy’s ability to identify cells and generate trajectories is enabled by the Python package Trackpy (Allan et al. 2019). Pandas, Numpy and Scipy are additional packages that provide TaxisPy with computational objects and functions required to store, filter and process the data generated by Trackpy. The python packages DEAP (Fortin et al. 2012) and Scoop (Hold-Geoffroy et al. 2014) are used to estimate parameter values for the identification of tumbles. All plotting routines are supported by Matplotlib.

**Figure 2.**
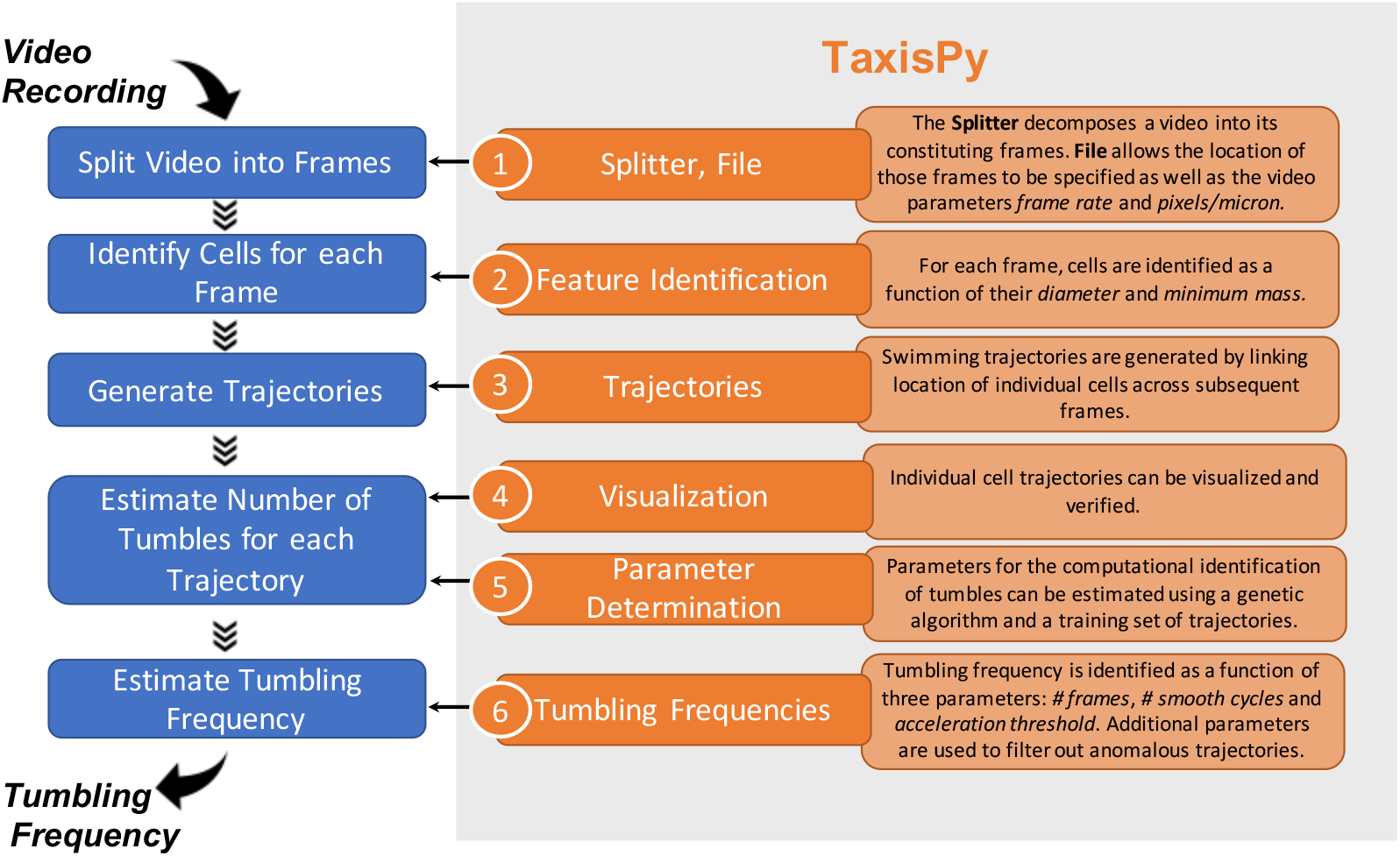
From a video recording to the bacterial tumbling frequency using TaxisPy. Five different tasks –represented by the blue rectangles– are involved in estimating the tumbling frequency of a bacterial culture from a video of its swimming behavior. TaxisPy’s user interface consists of six sequentially arranged tabs –represented by the orange rectangles– that provide required functionalities to perform each one of these tasks. Even though several parameters are involved in various steps of the analysis, TaxisPy provides necessary tools to identify appropriate values for those parameters.

1. Install Docker on your computer. Docker is supported by all major operating systems: Linux, MacOs and Windows.
2. Download the latest TaxisPy image by typing the following command on a Terminal (Mac, Linux) or Command Prompt window (Microsoft Windows):
`docker pull m1vg/taxispy`
3. Start a Docker container to access TaxisPy by typing the following command on the same window:
`docker run -d -p 8888:8888 m1vg/taxispy` That command will create a container without access to the files of the host computer (i.e., your computer). Files created within the container will be lost after the container is stopped. In order to grant access to files on the host computer, the previous command should be complemented with the flag `––mount:`
`docker run -d -p 8888:8888 --mount`
`type=bind,source=/Users,target=/Documents/host m1vg/taxispy`
Windows users should use:
`docker run -d -p 8888:8888 --mount`
`type=bind,source=//c/Users,target=/Documents/host`
`m1vg/taxispy`
4. Access TaxisPy by opening the following address on any internet browser (e.g., Google Chrome):
http://localhost:8888/

Quantitative characterization of the swimming behavior of a given bacterial culture by means of its tumbling frequency involves five consecutive steps, as shown in Fig. 2. First, a video registering freely swimming cells is split into its constituting frames. Each frame represents a snapshot of the culture at a given time point. Splitting can be done by the software FFmpeg (it supports a wide range of video formats), for which a simple user interface is provided within the TaxisPy Docker image; or by any other software designed for that purpose. Then, cells present in each frame are located and swimming trajectories identified by linking the position of individual cells through subsequent frames. Within TaxisPy, this functionality is provided by Trackpy 0.4.2 (Allan et al. 2019), a Python package that implements feature-finding and linking algorithms originally introduced by Crocker and Grier (1996). The next step consists of calculating the number of tumbles for each trajectory. Previous reports have used motion properties of individual trajectories – e.g., acceleration and velocities– to calculate the number of tumbles for each cellular trajectory. The underlying idea is that a cell decreases its linear velocity and increases its absolute acceleration below/above certain thresholds while tumbling. Specific threshold values were empirically determined by the authors of these studies (Berg and Brown 1972, Sager et al. 1988, Harwood 1989, Amsler 1996, Alon et al. 1998). It is clear that these values are a function of a variety of biological (i.e., bacterial strain, growth conditions, optical density, etc.) and technical (i.e., frame rate, signal smoothing) factors, which restricts the utility of such thresholds to the specific conditions for which they were estimated. TaxisPy addresses this issue by using a condition-specific training set consisting of a set of cellular trajectories with known changes of direction, along with a genetic algorithm. The generation of this training set by the user is supported by functionalities provided by TaxisPy. To estimate the number of tumbles for a given cellular trajectory, TaxisPy requires values for three different parameters: a threshold value for the absolute acceleration and values for two parameters controlling the smoothing routines: # Frames and # Smooth (see Table S1 in the Supplemental Information). Optimal values for these parameters are estimated based on a user-defined training set, so that the squared difference between observed and calculated number of tumbles by TaxisPy is minimized. TaxisPy uses the functionality provided by the Python package DEAP (Fortin et al. 2012) to solve this optimization problem. The final step in the characterization of the swimming behavior of a bacterial culture involves the determination of its average tumbling frequency. This value is calculated by dividing the total number of tumbles estimated for a set of trajectories over the total duration of those trajectories. Alternatively, the tumbling frequency can be calculated for each individual trajectory and an average over all trajectories can be used to estimate the average tumbling frequency of the culture. TaxisPy offers the flexibility to select the desired calculation method. In this work, reported tumbling frequencies were calculated using the first method, which divides the total number of tumbles over the total duration of all trajectories. In order to exclude anomalous trajectories that might bias the average tumbling frequency calculated for a set of trajectories, TaxisPy offers a series of filters to remove trajectories of nonmotile cells, trajectories exhibiting a too high number of tumbles (e.g., “stuck” cells that are trying to detach) or trajectories with a linear velocity that is abnormally low. Refer to the tutorial contained in the Docker image under **/Tutorial** for a more detailed description of this workflow.

### 2.4 Key parameter values required by TaxisPy

A comprehensive list of all parameters required by TaxisPy, along with a short description and nominal values is provided in Table S1 of the Supplemental Information. Eleven key parameters are involved in different stages of the analysis. These include parameters such as ***Diameter*** and ***Min. Mass***, which are used in the **Feature Identification** tab and are directly passed to Trackpy for the identification of individual cells in each frame. A nominal value of 25 pixels for ***Diameter*** and 2,000 for ***Min. Mass*** were used for all video analyses of both *P. putida* and *E. coli* cultures. TaxisPy offers an intuitive way to identify optimal values for these two parameters by visually inspecting the ability of the software to correctly identify cells in three different frames, corresponding to the first, middle, and last frame of a given set. Cells identified in these frames are enclosed by a blue circle by TaxisPy. Appropriate values for ***Diameter*** and ***Min. Mass*** will maximize the number of cells correctly identified by TaxisPy.

A second group of key parameters consists of ***# Trajectories***, ***Trajectory #*** and ***# Chng. Dir***. They refer to the number of trajectories within a given user-defined training set, the ID of each trajectory and the number of changes of direction or tumbles for each trajectory, respectively. These parameters define the training set that is used by DEAP to identify optimal values for smoothing parameters (***#Frames*** and ***#Smooth**)* and a value for the acceleration threshold (***Acc. Thrhld**.*), which are used to identify the number of tumbles of trajectories not contained within the training set. The genetic algorithm implemented by DEAP used a population size of 100 individuals and 5 generations. The initial population was generated at random, using bounds on ***Acc. Thrhld*** of 1000 micron/s^2^ and ***#Frames*** and ***#Smooth*** of 5. To increase variability within the population, we applied a two-point crossover and gaussian mutation with mean of 0 and standard deviation of 0.2. The independent probability for each attribute of the population to be mutated was 0.2. This configuration of the genetic algorithm exhibited a good performance for the training sets used in this study. Population size, number of generations and bounds of key parameters can be customized from the tab **Parameter Determination**. The training set used by DEAP can be easily generated by the user employing the tabs **Trajectories** and **Visualization**. Refer to Tables S2 to S6 in the Supplemental Information for training sets used to identify the acceleration threshold and smoothing parameters for the *P. putida* experiments and to Tables S8 to S10 for training sets used for the *E. coli* experiments. Optimal parameter values resulting from these training sets are summarized in Table S7 for the *P. putida* videos and in Table S11 for the *E. coli*.

An additional group of key parameters is located within the tab **Tumbling Frequencies**. These parameters control the way TaxisPy filters out anomalous trajectories and calculates average tumbling frequencies. The parameters ***Velocity***, ***Dsplcmt, %*** and ***Max. Chng. Dir**.* are used to sort out short trajectories, trajectories of cells swimming with too low linear velocities and trajectories exhibiting an excessive number of turns, respectively. Nominal values of 4 micron/s, 10%, and 10 tumbles were used to analyze all data presented in this study. Finally, the parameter ***T. Int. (s)*** is used to specify the time intervals used by TaxisPy to calculate the temporal evolution of the tumbling frequency of the culture. A value of 5 seconds was used for the analysis of the *P. putida* experiments, while a value of 10 seconds was used for *E. coli*.

## 3. Results

In order to demonstrate the utility of TaxisPy, the chemotactic response of two different microorganisms after stimulation with chemical attractants was video recorded and analyzed. Using the frames of the recorded videos as input, TaxisPy allowed the quantification of the effect of shikimate on the tumbling frequency of *P. putida* and the visualization of the adaptation process of *E. coli* after exposure to 1 mM L-aspartate. The following results were generated using parameter values listed in Table S7 for *P. putida* F1 and Table S11 for *E. coli*. Refer to ReadMe Notebooks contained in the Docker image under **/Ecoli_Aspartate** and **/Pseudomonas_Shikimate** to reproduce individual results.

### 3.1 Characterization of the chemotactic response of *P. putida*

*P. putida* F1 was exposed to four different concentrations of shikimate. Chemotaxis buffer without shikimate was used as the control condition. The response of the culture under each condition was video recorded in triplicate for sixty seconds and TaxisPy was subsequently used to calculate the average tumbling frequency under each condition as described in the Methods section. Fig. 3 summarizes our findings. As expected, the cellular tumbling frequency decreases from its basal value of 0.42 tumbles/second under unstimulated conditions to a low value of 0.023 tumbles/second for a shikimate concentration of 1 mM.

**Figure 3.**
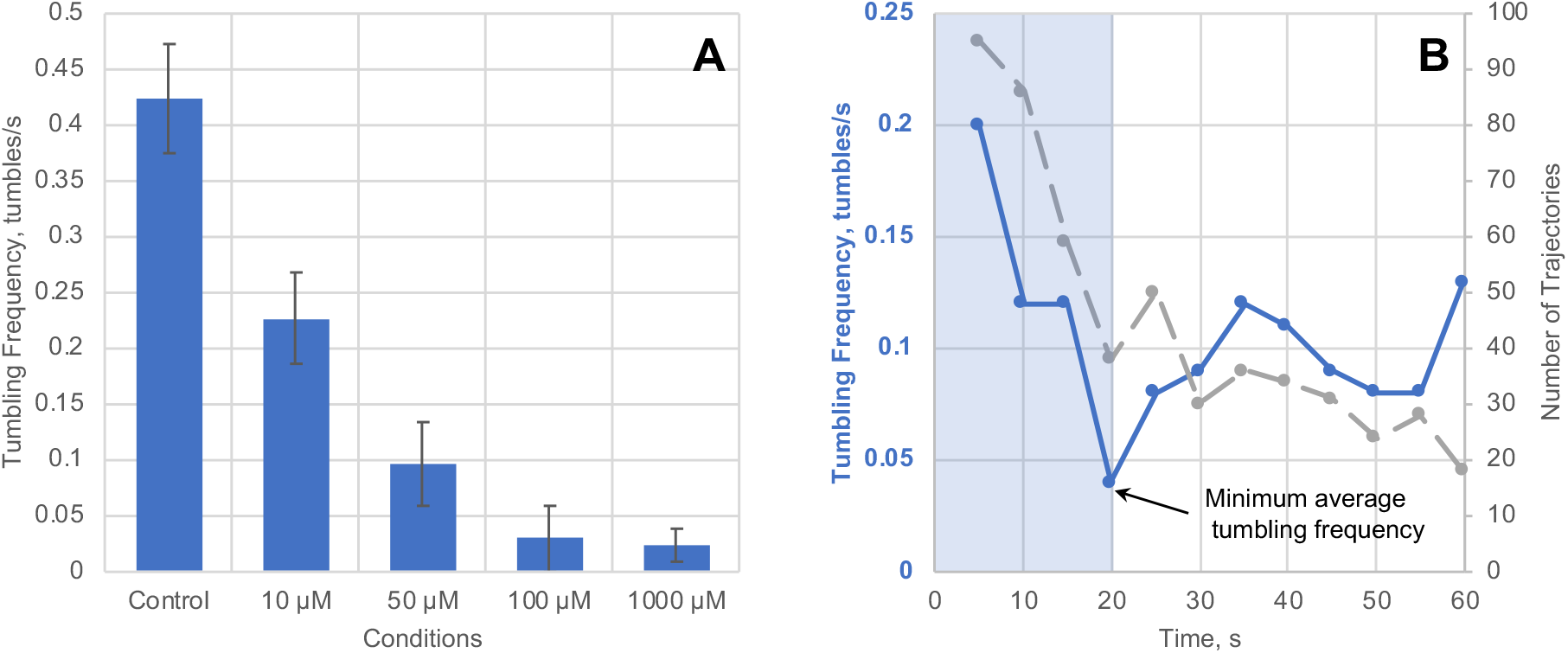
Tumbling frequency of *P. putida* for different shikimate concentrations. **A.** Tumbling frequency exhibited by the culture as a function of the shikimate concentration. After stimulation with shikimate, the swimming pattern was characterized by the minimum average tumbling frequency exhibited by the cells during the first 20 seconds and calculated using time intervals of 5 seconds. For the control cultures (chemotaxis buffer only), the swimming pattern was characterized by the average tumbling frequency calculated for the first 20 seconds of each video. Refer to main text for details. The number of trajectories used to calculate tumbling frequencies for each video ranged from 257 to 23. Error bars represent one standard deviation of the culture’s tumbling frequency calculated from three different videos. **B.** Temporal evolution of the tumbling frequency of *P. putida* after exposure to 1 mM shikimate is represented by the blue solid line. Average frequencies were calculated using intervals of 5 seconds; the corresponding data point is placed at the end of each interval. The grey dashed line represents the number of cellular trajectories used by TaxisPy to calculate reported average frequencies. The shaded blue area represents the first 20 seconds of the video used to determine the minimum average tumbling frequency.

The *minimum* tumbling frequency exhibited by the culture within the first 20 seconds of each video was used in Fig. 3A to assess the effect of different concentrations of shikimate on the swimming pattern of the culture. This value was used instead of the *average* tumbling frequency over the same period of time because it better captured the reduction of the culture’s tumbling frequency after exposure to the chemical attractant. This is justified in Fig. 3B, where the temporal evolution of the tumbling frequency after addition of 1 mM of shikimate is shown. While the average tumbling frequency for the first 20 seconds corresponds to 0.13 tumbles/second, the minimum tumbling frequency within the same period of time corresponds to 0.04 tumbles/second. The origin of the discrepancy between these two values can be attributed to temporal processes involved in the diffusion of the chemical attractant in the culture’s medium, as well as to temporal processes involved in cellular sensing and response to the chemical gradient. Note that the analysis was restricted to the initial 20 seconds of the video instead of its total duration (see blue rectangle in Fig. 3B). There are two reasons for this. The first one is related to the adaptation process that the cellular sensing machinery undergoes and the associated increase in the tumbling frequency. The second reason is related to the steady decrease over time in the number of cellular trajectories identified by TaxisPy, which compromises the representativeness of the tumbling frequencies calculated and potentially increases the noise at later time points. This observation was consistent over all recorded videos (see Fig. S1). For consistency, the tumbling frequency for the control condition was also calculated over a period of 20 seconds.

### 3.2 Visualizing the adaptation process of the chemotactic machinery of *E. coli*

The two-component system controlling the chemotactic response of *E. coli* is able to adapt to external stimuli (Macnab & Koshland 1972, Berg & Tedesco 1975, Alon et al. 1999). Adaptation refers to the temporal process by which the cellular tumbling frequency returns to its pre-stimulation state. As a proof-of-concept and to demonstrate the utility and versatility of TaxisPy in elucidating this process, we stimulated an *E. coli* culture with 1 mM L-aspartate and videotaped its swimming pattern at regular time intervals for 40 minutes. Each video was split into its constituting frames and these were analyzed by TaxisPy. Tumbling frequencies were calculated using time intervals of 10 seconds. These data are graphically represented by the blue dots in Fig. 4. Black dots in the same figure represent the tumbling frequency of a control culture without the addition of the chemical attractant.

**Figure 4.**
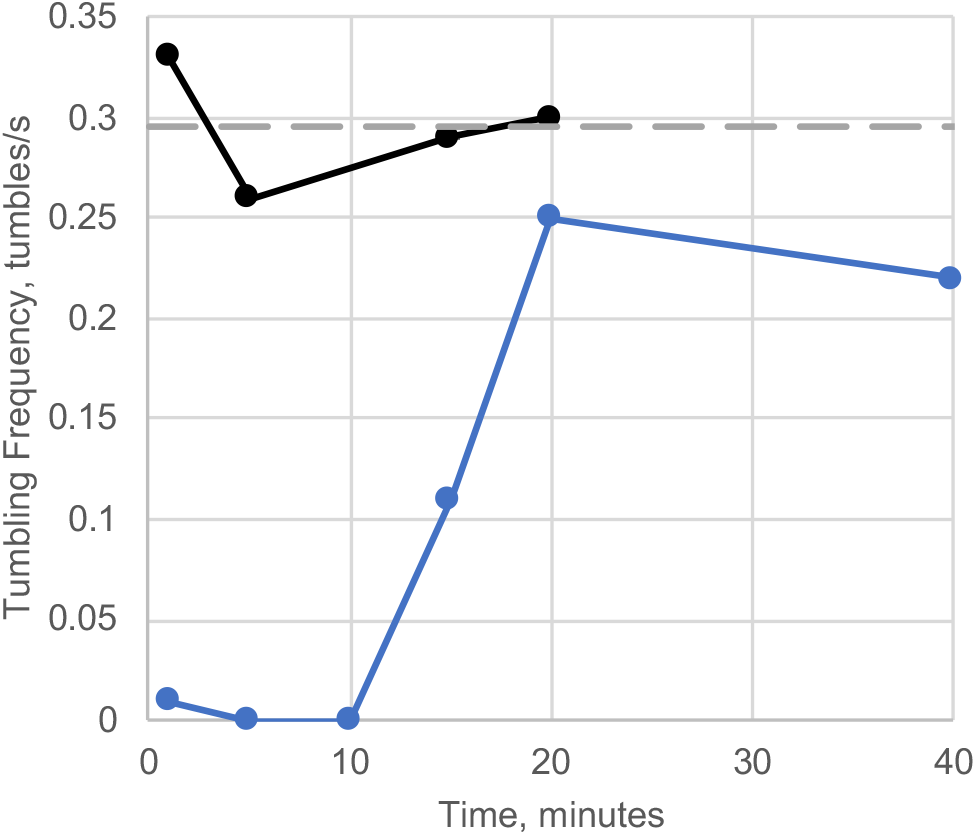
Adaptation of the chemotactic machinery of *E. coli*. L-aspartate (1 mM) was added to an *E. coli* culture at time 0 and its swimming pattern video tapped for 10 seconds in regular time intervals over 40 minutes. Each video was subsequently analyzed using TaxisPy to estimate the temporal evolution of the tumbling frequency. These data points are represented by the blue dots. The tumbling frequency of a control culture without attractant is represented by the black dots. The grey dashed line represents the average tumbling frequency of the unstimulated culture (0.295 tumbles/second). Each data point was generated from the analysis of 86 to 420 individual cellular trajectories.

The expected adaptation behavior is evident from the time course of the tumbling frequency for the stimulated culture (blue solid line in Fig. 4). As time goes by, the average tumbling frequency of the culture increases from values lower than 0.05 tumbles/second during the first 10 minutes, to reach a maximum value of 0.25 tumbles/second after 20 minutes. The adaptation process can be characterized by two parameters: the precision of adaptation and the adaptation time (Alon et al. 1999). The first parameter is defined as the ratio between the average tumbling frequency of the *unstimulated* culture and the average tumbling frequency of the *stimulated* culture after adaptation has been reached. The second parameter is defined as the time where the tumbling frequency of stimulated cells rises to halfway between its earliest measured value and its steady-state value. For the specific system under analysis, the adaptation time was 15 minutes and the precision of adaptation was 1.18.

## 4. Discussion

A major difficulty related to the customized use of cell tracking software is the determination of key parameter values for the identification of tumbles or changes of direction. In the past, such parameters were empirically determined (Berg and Brown 1972, Harwood et al. 1989), which compromises the applicability of those parameters under different conditions – e.g., different microorganisms or stimuli. TaxisPy systematically addresses this issue by implementing various strategies designed to guide the selection of relevant parameter values. For instance, TaxisPy employs a genetic algorithm along with a condition-specific training set to identify optimal values for three key parameters: *# Frames*, *# Smooth* and *Acc. Threshold.* The first two parameters affect the way that TaxisPy smooth cellular velocity data by calculating the average from a number of frames (dictated by *# Frames*) a certain number of times (determined by *# Smooth*). Noisy positional data tend to require more smoothing cycles. TaxisPy analyzes the time course of the absolute acceleration –which is calculated from the smoothed velocity data– to identify tumbling events. We use the premise that a cell decreases its linear velocity when tumbling and increases it again thereafter. Thus, a change of direction or tumble is characterized by two adjacent peaks in the time course of the absolute acceleration. To avoid identifying normal fluctuations in the velocity during smooth swimming as a tumble, the acceleration peaks are required to surpass a threshold value dictated by *Acc. Threshold*. Since smoothing the velocity data decreases the height of the peaks in the acceleration plot, high values for *# Frames* and *# Smooth* are usually accompanied by low values for *Acc. Threshold* and vice versa.

One of the motivations for the development of TaxisPy was to offer a simple method for the quantitative analysis of bacterial chemotaxis. Following the way paved by previous studies that implemented video-based analysis of bacterial chemotaxis, we decided to base our method on the analysis of cellular motion data for the identification of tumbling frequencies. This led us to identify the necessity of condition-specific values for some parameters –*# Frames*, *# Smooth* and *Acc. Threshold*– and to develop features in TaxisPy for their identification. Even though constructing training sets and identifying optimal parameter values usually requires a couple of minutes, this procedure can become tedious if a large number of videos need to be processed. A promising approach for the “parameter-free” identification of tumbles is the analysis of cellular trajectories by multilayer neural networks (Lecun et al. 1998, Ciregan et al. 2012). These networks could be first trained on a large data set of cellular trajectories –each one of them labeled with the number of tumbles– to then be used on new trajectories to estimate their number of tumbles. An advantage of such an approach would be that the model parameterization would be done just once and could be then applied for various microorganisms under different conditions. Paving the way towards this new approach, we are providing an initial training set in the Supplementary File 1, which was manually extracted and curated from the *P. putida* experiments and consists of over 700 cellular trajectories, each one provided with the number of observed tumbles.

In this study, we used two different experimental set-ups to study the bacterial chemotactic response. For *P. putida*, the temporal response to shikimate pulses of varying concentration were measured on *microscope glass slides*. Using the minimum tumbling frequency during the first 20 seconds of each experiment, we were able to quantitatively capture the effect of the shikimate concentration on the swimming behavior. As expected, we observed a decrease in the tumbling frequency as the concentration of shikimate was increased (Fig. 3A). Increasing the shikimate concentration over 100 µM did not seem to have a further effect on the chemotactic response of *P. putida*. Additionally, values calculated for the tumbling frequency were within the same order of magnitude as previous reports (Harwood et. al. 1989), which validates the computations performed by TaxisPy.

As mentioned in the results section and shown in Figs. 3B and S1, the number of trajectories identified by TaxisPy consistently decreased during the course of each experiment. This is potentially related to a temperature or light-dependent taxis away from the observation field of the microscope. An indirect support to this claim is provided by a report by Paster and Ryu (2008), in which a tactic response was observed in *E. coli* as a function of temperature gradients. The authors showed that at temperatures below 31 °C, the response to thermal stimuli is similar to the chemotactic response. However, at temperatures above 31°C, some cells showed an inverted response, switching from warm- to cold-seeking behavior. Additional factors to be considered might be related with oxygen depletion and the accompanied overall loss of bacterial motility (Douarche et al. 2009) and with cellular migration towards the nearest air/water interface due to aerotaxis (Taylor 1983) and away from the focus of the microscope. From a practical point of view, a steady decrease in the number of cellular trajectories over time has two implications. First, it suggests that the uncertainty of data extracted from adaptation experiments performed on glass slides will increase over time, since the number of trajectories used to calculate tumbling frequencies will continuously decrease. Second, it poses a biological constraint on the maximal length of the videos when the swimming pattern is observed on a glass slide. In line with these observations, we limited the analysis of each *P. putida* video to the first 20 seconds.

In the case of *E. coli*, a different experimental set-up was followed. Since the adaptation time exhibited by this microorganism under the studied conditions is in the order of magnitude of minutes, the actual adaptation process was conducted in an *Eppendorf tube* instead of in a microscope slide. Samples were taken from the vessel, which was kept under shaking conditions, and analyzed under the microscope for 10 seconds. In this way, we were able to estimate both the adaptation time and the precision of adaptation of the culture. The estimated value for the adaptation time of 15 minutes and the precision of adaptation of 1.18 are consistent with previously reported values (Alon et. al. 1999). These results serve as further validation of our methods.

TaxisPy was designed for users with limited programming knowledge. Its user interface provides necessary tools to split a video into its constituting frames, identify cells in individual frames, link these to obtain trajectories, filter anomalous trajectories and calculate tumbling frequencies. Thanks to its convenient distribution through Docker, intuitive user interface and biologically feasible results, TaxisPy represents a valuable computational tool for the quantitative analysis of bacterial chemotaxis.

## 5. Acknowledgements

This work was supported by grants from the National Science Foundation to R.E.P and M.A.S (MCB 1716833), and the USDA National Institute of Food and Agriculture, Hatch/Evans-Allen/McIntire Stennis project 1020219 to R.E.P.

## 6. Author Contributions

Conceptualization, R.E.P.; Methodology, R.A.S. and M.A.V.; Software, M.A.V; Validation, R.A.S and M.A.V.; Writing, M.A.V., R.A.S. and R.E.P; Supervision, R.E.P. and M.A.S., Funding Acquisition, R.E.P and M.A.S.

