## Supplemental Information for "TaxisPy: A Python-based Software for the Quantitative Analysis of Bacterial Chemotaxis"

**Table S1**. **Parameters involved in a quantitative chemotaxis analysis by TaxisPy.** Parameters required by the software in each one of its different windows are listed below, as well as a short description and their nominal values. Even though the number of parameters appears to be large, TaxisPy provides intuitive tools designed to guide the selection of specific values. Key parameters are marked by a blue asterisk (*).

| Tab | Parameter | Description | Nominal Value |
|---|---|---|---|
| **File** | *Path* | Indicates location of folder containing frames | -- |
| | *Frames* | Indicates range of frames to be analyzed | 0 - 410 frames for *P. putida* videos; 0 - 202 frames for *E. coli* videos. |
| | *Frames/s* | Frame rate of video. Corresponds to the ratio between the total number of frames of a video and its duration. | 20.3 frames/s |
| | *Pixels/Micron* | It is an indication of the resolution of the video camera. Can be determined by taking a snapshot of a scale and counting the number of pixels contained in one micron. | 6 pixels/micron |
| **Feature Identification** | *Diameter** | Diameter of an average cell in pixels. It should be an odd integer. | 25 pixels |
| | *Min. Mass** | The minimum integrated brightness. This is a crucial parameter for eliminating spurious features. | 2000 |
| | *Invert?* | The software looks for bright features. Set to true if cells are dark objects. | TRUE |
| **Trajectories** | *Max. Disp.* | The maximum distance cells can move between frames | 25 pixels |
| | *Min. # Frms* | Minimum number of frames for a trajectory to be considered by the software | 20 frames |
| | *Disp. Thrshld* | Minimum length in % that a trajectory should have to be shown. A value of 0% shows all trajectories, while 100% shows the longest trajectory. | 0% |

| Tab | Parameter | Description | Nominal Value |
|---|---|---|---|
| | *Frame Rng* | Controls the frame range for which trajectories are shown. | -- |
| **Visualization** | *Trajectory* | Trajectory to be shown in a two-dimensional plot and in video. | Last trajectory |
| | *Frame #* | Controls the frame that is shown | 0 |
| **Parameter Determination (Analysis of Individual Trajectories)** | *Trajectory #* | Trajectory for which motion analysis is performed | Last trajectory |
| | *Time Range* | Time range in which motion analysis is performed | tin – tf. These variables correspond to initial and final time points of the trajectory, respectively. |
| | *# Frames* | Number of subsequent frames used to calculate an average velocity | 4 frames |
| | *# Smooth* | Number of times (cycles) that the average velocity is calculated | 3 cycles |
| | *Acc. Thrhld* | Acceleration threshold that a trajectory should surpass for the software to identify a tumble. Absolute acceleration values are considered. | 10 micron/s/s |
| **Parameter Determination (Genetic Algorithm)** | *# Trajectories** | Number of trajectories in the training set | 10 |
| | *Trajectory #** | ID of a given trajectory within the training set | random |
| | *# Chng. Dir.** | Number of tumbles or changes of direction exhibited by a given trajectory within the training set | 0 |
| **Tumbling Frequencies** | *# Frames** | Number of subsequent frames used to calculate an average velocity | 4 frames |
| | *# Smooth** | Number of times (cycles) that the average velocity is calculated | 3 cycles |
| | *Acceleration** | Acceleration threshold that a trajectory should surpass for the software to identify a tumble. Absolute acceleration values are considered. | 10 micron/s/s |
| | *Velocity** | Minimum velocity that a cell should exhibit to be considered in the calculation of the tumbling frequency | 4 micron/s |
| | *Dsplcmt, %** | Minimum displacement (in %) that a trajectory should exhibit to be considered in the calculation of the tumbling frequency | 10% |
| | *Max. Chng Dir** | Maximum number of tumbles that a trajectory can exhibit to be considered in the calculation of the tumbling frequency. Trajectories with a higher number of tumbles are ignored. | 10 |
| | *Frame Rng* | Frame range for which the tumbling frequency is calculated. | 0 - 410 frames for *P. putida* videos; 0 - 202 frames for *E. coli* videos. |

| Tab | Parameter | Description | Nominal Value |
|---|---|---|---|
| | *T. Int. (s)** | Time interval used to calculate tumbling frequencies | 1 second |
| | *Chg. Dir. (1/s)* | Tumbling frequency used to estimate adaptation time. | 0.45 tumbles/second |

**Table S2. Training sets stemming from control cultures of *P. putida*.** Training sets were used to identify optimal values for the acceleration threshold as well as for smoothing parameters –*# Frames* and *# Smooth*–, which are reported in Table S7. Training sets for each video were generated using tabs **Trajectories** and **Visualization** of TaxisPy.

| **Control** | | | | | | | | |
|---|---|---|---|---|---|---|---|---|
| **CB0** | | | **CB1** | | | **CB2** | | |
| # | Trajectory ID | Tumbles | # | Trajectory ID | Tumbles | # | Trajectory ID | Tumbles |
| 1 | 125 | 1 | 1 | 364 | 2 | 1 | 186 | 1 |
| 2 | 190 | 0 | 2 | 526 | 0 | 2 | 290 | 0 |
| 3 | 167 | 1 | 3 | 555 | 1 | 3 | 190 | 1 |
| 4 | 145 | 1 | 4 | 364 | 2 | 4 | 42 | 1 |
| 5 | 1229 | 1 | 5 | 778 | 0 | 5 | 214 | 1 |
| 6 | 1032 | 1 | 6 | 891 | 1 | 6 | 517 | 1 |
| 7 | 991 | 1 | 7 | 600 | 1 | 7 | 859 | 1 |
| 8 | 1433 | 1 | 8 | 1166 | 1 | 8 | 874 | 1 |
| 9 | 1045 | 0 | 9 | 1233 | 1 | 9 | 702 | 0 |
| 10 | 1415 | 1 | 10 | 1259 | 0 | 10 | 1240 | 0 |
| 11 | 559 | 1 | 11 | 42 | 2 | 11 | 1108 | 2 |
| 12 | 512 | 1 | 12 | 600 | 1 | 12 | 1262 | 4 |
| 13 | 1015 | 3 | 13 | 332 | 2 | 13 | 1240 | 0 |
| 14 | 326 | 2 | 14 | 546 | 1 | 14 | 186 | 1 |
| | | | 15 | 433 | 1 | 15 | 1105 | 1 |
| | | | 16 | 318 | 1 | 16 | 984 | 1 |
| | | | 17 | 650 | 0 | 17 | 832 | 3 |

**Table S3. Training sets stemming from stimulation experiments of *P. putida* cultures with 10 µM shikimate.** Training sets were used to identify optimal values for the acceleration threshold as well as for smoothing parameters –# *Frames* and # *Smooth*–, which are reported in Table S7. Training sets for each video were generated using tabs **Trajectories** and **Visualization** of TaxisPy.

| **10 µM shikimate** | | | | | | | | |
|---|---|---|---|---|---|---|---|---|
| **shik9** | | | **shik10** | | | **shik11** | | |
| # | Trajectory ID | Tumbles | # | Trajectory ID | Tumbles | # | Trajectory ID | Tumbles |
| 1 | 22 | 1 | 1 | 208 | 1 | 1 | 299 | 1 |
| 2 | 624 | 0 | 2 | 396 | 0 | 2 | 220 | 1 |
| 3 | 600 | 1 | 3 | 372 | 0 | 3 | 38 | 1 |
| 4 | 638 | 1 | 4 | 49 | 1 | 4 | 160 | 0 |
| 5 | 665 | 0 | 5 | 194 | 0 | 5 | 127 | 2 |
| 6 | 184 | 1 | 6 | 413 | 0 | 6 | 500 | 1 |
| 7 | 1131 | 1 | 7 | 324 | 2 | 7 | 59 | 1 |
| 8 | 1286 | 1 | 8 | 649 | 1 | 8 | 822 | 1 |
| 9 | 1026 | 2 | 9 | 441 | 1 | 9 | 709 | 1 |
| 10 | 1840 | 1 | 10 | 762 | 1 | 10 | 546 | 1 |
| 11 | 1680 | 1 | 11 | 637 | Ta1 | 11 | 1047 | 0 |
| 12 | 1712 | 0 | 12 | 923 | 0 | 12 | 1035 | 0 |
| 13 | 1646 | 1 | 13 | 885 | 1 | 13 | 999 | 1 |
| | | | 14 | 869 | 1 | | | |

**Table S4. Training sets stemming from stimulation experiments of *P. putida* cultures with 50 µM shikimate.** Training sets were used to identify optimal values for the acceleration threshold as well as for smoothing parameters –# *Frames* and # *Smooth*–, which are reported in Table S7. Training sets for each video were generated using tabs **Trajectories** and **Visualization** of TaxisPy.

| 50 µM shikimate | | | | | | | | |
|---|---|---|---|---|---|---|---|---|
| shik12 | | | shik13 | | | shik14 | | |
| # | Trajectory ID | Tumbles | # | Trajectory ID | Tumbles | # | Trajectory ID | Tumbles |
| 1 | 499 | 1 | 1 | 1167 | 0 | 1 | 717 | 3 |
| 2 | 708 | 1 | 2 | 203 | 0 | 2 | 25 | 0 |
| 3 | 149 | 0 | 3 | 26 | 1 | 3 | 1083 | 1 |
| 4 | 110 | 0 | 4 | 621 | 0 | 4 | 280 | 1 |
| 5 | 472 | 1 | 5 | 81 | 1 | 5 | 979 | 1 |
| 6 | 1233 | 1 | 6 | 250 | 1 | 6 | 930 | 0 |
| 7 | 1302 | 1 | 7 | 21 | 1 | 7 | 1274 | 1 |
| 8 | 1153 | 1 | 8 | 780 | 1 | 8 | 1440 | 0 |
| 9 | 899 | 1 | 9 | 609 | 1 | 9 | 1119 | 2 |
| 10 | 1001 | 1 | 10 | 775 | 0 | 10 | 1810 | 2 |
| 11 | 1700 | 1 | 11 | 666 | 1 | 11 | 1664 | 2 |
| 12 | 2122 | 1 | 12 | 878 | 1 | 12 | 1942 | 0 |
| 13 | 2239 | 2 | 13 | 868 | 1 | 13 | 1700 | 1 |
| 14 | 1639 | 0 | 14 | 494 | 0 | 14 | 2345 | 0 |
| | | | 15 | 1090 | 1 | 15 | 2573 | 0 |
| | | | 16 | 961 | 0 | 16 | 2352 | 0 |
| | | | 17 | 149 | 0 | 17 | 2356 | 0 |
| | | | 18 | 112 | 0 | 18 | 2394 | 0 |
| | | | 19 | 204 | 0 | 19 | 2449 | 0 |
| | | | 20 | 203 | 0 | 20 | 2398 | 0 |

**Table S5. Training sets stemming from stimulation experiments of *P. putida* cultures with 100 µM shikimate.** Training sets were used to identify optimal values for the acceleration threshold as well as for smoothing parameters –*# Frames* and *# Smooth*–, which are reported in Table S7. Training sets for each video were generated using tabs **Trajectories** and **Visualization** of TaxisPy.

| 100 µM shikimate | | | | | | | | |
|---|---|---|---|---|---|---|---|---|
| **shik6** | | | **shik7** | | | **shik8** | | |
| # | Trajectory ID | Tumbles | # | Trajectory ID | Tumbles | # | Trajectory ID | Tumbles |
| 1 | 314 | 0 | 1 | 143 | 0 | 1 | 42 | 1 |
| 2 | 167 | 0 | 2 | 201 | 0 | 2 | 217 | 0 |
| 3 | 628 | 0 | 3 | 589 | 0 | 3 | 123 | 1 |
| 4 | 16 | 1 | 4 | 876 | 0 | 4 | 180 | 0 |
| 5 | 247 | 0 | 5 | 2141 | 0 | 5 | 50 | 1 |
| 6 | 220 | 0 | 6 | 1920 | 0 | 6 | 90 | 0 |
| 7 | 964 | 0 | 7 | 1274 | 0 | 7 | 355 | 0 |
| 8 | 933 | 0 | 8 | 3650 | 0 | 8 | 430 | 0 |
| 9 | 703 | 1 | 9 | 2821 | 0 | 9 | 372 | 0 |
| 10 | 1122 | 0 | 10 | 3363 | 0 | 10 | 408 | 0 |
| 11 | 1169 | 0 | 11 | 2790 | 1 | 11 | 582 | 0 |
| 12 | 1120 | 0 | 12 | 3175 | 0 | 12 | 627 | 0 |
| 13 | 1180 | 0 | 13 | 3076 | 0 | 13 | 691 | 0 |
| | | | | | | 14 | 514 | 0 |

**Table S6. Training sets stemming from stimulation experiments of *P. putida* cultures with 1000 µM shikimate.** Training sets were used to identify optimal values for the acceleration threshold as well as for smoothing parameters –# *Frames* and # *Smooth*–, which are reported in Table S7. Training sets for each video were generated using tabs **Trajectories** and **Visualization** of TaxisPy.

| **1000 µM shikimate** | | | | | | | | |
|---|---|---|---|---|---|---|---|---|
| **shik3** | | | **shik4** | | | **shik5** | | |
| # | Trajectory ID | Tumbles | # | Trajectory ID | Tumbles | # | Trajectory ID | Tumbles |
| 1 | 135 | 0 | 1 | 104 | 0 | 1 | 326 | 0 |
| 2 | 151 | 0 | 2 | 7 | 0 | 2 | 392 | 0 |
| 3 | 570 | 1 | 3 | 143 | 0 | 3 | 221 | 0 |
| 4 | 121 | 1 | 4 | 257 | 0 | 4 | 16 | 0 |
| 5 | 572 | 1 | 5 | 274 | 1 | 5 | 529 | 0 |
| 6 | 105 | 1 | 6 | 263 | 0 | 6 | 523 | 0 |
| 7 | 812 | 0 | 7 | 750 | 0 | 7 | 748 | 0 |
| 8 | 979 | 1 | 8 | 947 | 0 | 8 | 592 | 0 |
| 9 | 1257 | 0 | 9 | 892 | 0 | 9 | 929 | 0 |
| 10 | 1179 | 1 | 10 | 1007 | 0 | 10 | 1205 | 0 |
| 11 | 1562 | 0 | 11 | 1548 | 0 | 11 | 919 | 0 |
| 12 | 1462 | 0 | 12 | 1278 | 0 | 12 | 1105 | 0 |
| 13 | 1424 | 0 | 13 | 1259 | 0 | 13 | 914 | 0 |

**Table S7. Key parameter values used in the analysis of *P. putida* stimulation experiments**. The condition-specific parameter values listed below –#Frames, #Smooth and Acc. Threshold– were automatically determined using a genetic algorithm and corresponding training sets (Tables S2 to S6). Values for both Tumbling Freq. and Min. Tumbling Freq. were calculated employing those condition-specific parameter values and the tab **Tumbling Frequencies** of TaxisPy. *Tumbling Freq*. refers to the culture's tumbling frequency during the first 20 seconds. On the other hand, *Min. Tumbling Freq*. refers to the minimum tumbling frequency exhibited by the culture during the same period of time. Time intervals of 5 seconds were used to calculate the temporal evolution of the tumbling frequency in that case. Note that columns *Nr. Traj.* contain the number of trajectories used to calculate either *Tumbling Freq*. or *Min. Tumbling Freq*.

| Condition | Video ID | # Frames | # Smooth | Acc. Threshold | Tumbling Freq. | Nr. Traj. | Min. Tumbling Freq. | Nr. Traj. |
|---|---|---|---|---|---|---|---|---|
| **Control** | CB0 | 2 | 4 | 167 | 0.46 | 257 | -- | -- |
| | CB1 | 3 | 4 | 67 | 0.36 | 163 | -- | -- |
| | CB2 | 2 | 5 | 100 | 0.43 | 222 | -- | -- |
| **10 μM shikimate** | shik9 | 4 | 3 | 60 | 0.26 | 313 | 0.24 | 116 |
| | shik10 | 2 | 5 | 103 | 0.33 | 119 | 0.26 | 48 |
| | shik11 | 2 | 3 | 140 | 0.26 | 162 | 0.18 | 42 |
| **50 μM shikimate** | shik12 | 5 | 4 | 80 | 0.07 | 263 | 0.07 | 71 |
| | shik13 | 5 | 2 | 103 | 0.16 | 151 | 0.08 | 38 |
| | shik14 | 2 | 5 | 140 | 0.27 | 413 | 0.14 | 73 |
| **100 μM shikimate** | shik6 | 4 | 4 | 71 | 0.07 | 233 | 0.06 | 52 |
| | shik7 | 2 | 4 | 160 | 0.06 | 142 | 0.03 | 35 |
| | shik8 | 5 | 5 | 140 | 0 | 109 | 0 | 23 |
| **1000 μM shikimate** | shik3 | 4 | 3 | 80 | 0.13 | 278 | 0.04 | 38 |
| | shik4 | 5 | 4 | 60 | 0.05 | 133 | 0.02 | 46 |
| | shik5 | 3 | 5 | 120 | 0.03 | 149 | 0.01 | 33 |

**Table S8**. **Training sets stemming from control cultures of *E. coli.*** Training sets were used to identify optimal values for the acceleration threshold as well as for smoothing parameters –# *Frames* and # *Smooth*–, which are reported in Table S11. Training sets for each video were generated using tabs **Trajectories** and **Visualization** of TaxisPy.

| **Control** | | | | | | | | | | | |
|---|---|---|---|---|---|---|---|---|---|---|---|
| **RPMSB_2 @ 1 min** | | | **RPMSB_3 @ 5 min** | | | **RPMSB_4 @ 15 min** | | | **RPMSB_5 @ 20 min** | | |
| # | ID | Tumbles | # | ID | Tumbles | # | ID | Tumbles | # | ID | Tumbles |
| 1 | 489 | 2 | 1 | 69 | 1 | 1 | 236 | 1 | 1 | 303 | 1 |
| 2 | 203 | 2 | 2 | 306 | 0 | 2 | 75 | 2 | 2 | 705 | 0 |
| 3 | 123 | 0 | 3 | 124 | 1 | 3 | 315 | 0 | 3 | 760 | 0 |
| 4 | 227 | 2 | 4 | 319 | 0 | 4 | 84 | 2 | 4 | 346 | 1 |
| 5 | 55 | 2 | 5 | 165 | 2 | 5 | 7 | 0 | 5 | 1178 | 1 |
| 6 | 912 | 2 | 6 | 427 | 2 | 6 | 739 | 1 | 6 | 1331 | 0 |
| 7 | 681 | 1 | 7 | 556 | 2 | 7 | 677 | 1 | 7 | 845 | 1 |
| 8 | 880 | 2 | 8 | 480 | 0 | 8 | 497 | 2 | 8 | 990 | 1 |
| 9 | 966 | 0 | 9 | 492 | 1 | 9 | 831 | 0 | 9 | 1045 | 1 |
| 10 | 636 | 0 | 10 | 435 | 2 | 10 | 485 | 0 | 10 | 1305 | 1 |
| 11 | 1160 | 0 | 11 | 449 | 0 | 11 | 917 | 0 | 11 | 1653 | 1 |
| 12 | 1037 | 1 | 12 | 626 | 0 | 12 | 864 | 1 | 12 | 1743 | 0 |
| 13 | 1251 | 0 | 13 | 681 | 1 | 13 | 943 | 1 | 13 | 1645 | 1 |
| 14 | 1270 | 0 | 14 | 614 | 1 | 14 | 446 | 0 | 14 | 1497 | 0 |
| 15 | 1031 | 1 | 15 | 843 | 0 | 15 | 172 | 1 | 15 | 1591 | 0 |

**Table S9. Training sets stemming from stimulated cultures of *P. putida* during the first 10 minutes of adaptation.** Training sets were used to identify optimal values for the acceleration threshold as well as for smoothing parameters –# *Frames* and # *Smooth*–, which are reported in Table S11. Training sets for each video were generated using tabs **Trajectories** and **Visualization** of TaxisPy.

| **1 min - 10 Min** | | | | | | | | |
|---|---|---|---|---|---|---|---|---|
| **RPMSB_26 @ 1 min** | | | **RPMSB_27 @ 5 min** | | | **RPMSB_42 @ 10 min** | | |
| # | Trajectory ID | Tumbles | # | Trajectory ID | Tumbles | # | Trajectory ID | Tumbles |
| 1 | 454 | 0 | 1 | 64 | 0 | 1 | 325 | 0 |
| 2 | 310 | 0 | 2 | 217 | 0 | 2 | 50 | 0 |
| 3 | 3 | 1 | 3 | 290 | 0 | 3 | 214 | 0 |
| 4 | 392 | 0 | 4 | 270 | 0 | 4 | 207 | 0 |
| 5 | 15 | 0 | 5 | 226 | 0 | 5 | 149 | 0 |
| 6 | 776 | 0 | 6 | 512 | 0 | 6 | 319 | 0 |
| 7 | 589 | 0 | 7 | 493 | 0 | 7 | 356 | 0 |
| 8 | 732 | 0 | 8 | 737 | 0 | 8 | 474 | 0 |
| 9 | 881 | 0 | 9 | 707 | 0 | 9 | 552 | 0 |
| 10 | 753 | 0 | 10 | 828 | 0 | 10 | 350 | 0 |
| 11 | 926 | 0 | 11 | 911 | 0 | 11 | 504 | 0 |
| 12 | 924 | 0 | 12 | 1043 | 0 | 12 | 539 | 0 |
| 13 | 916 | 0 | 13 | 899 | 0 | 13 | 690 | 0 |
| 14 | 1002 | 0 | 14 | 1062 | 0 | 14 | 702 | 0 |
| 15 | 1030 | 0 | 15 | 1028 | 0 | 15 | 675 | 0 |

**Table S10. Training sets stemming from stimulated cultures of *P. putida* after 15, 20 and 40 minutes of adaptation.** Training sets were used to identify optimal values for the acceleration threshold as well as for smoothing parameters –# *Frames* and # *Smooth*–, which are reported in Table S11. Training sets for each video were generated using tabs **Trajectories** and **Visualization** of TaxisPy.

| **15 min - 40 Min** | | | | | | | | |
|---|---|---|---|---|---|---|---|---|
| **RPMSB_43 @ 15 min** | | | **RPMSB_30 @ 20 min** | | | **RPMSB_45 @ 40 min** | | |
| # | Trajectory ID | Tumbles | # | Trajectory ID | Tumbles | # | Trajectory ID | Tumbles |
| 1 | 112 | 0 | 1 | 198 | 1 | 1 | 281 | 0 |
| 2 | 102 | 0 | 2 | 335 | 1 | 2 | 11 | 2 |
| 3 | 163 | 0 | 3 | 91 | 2 | 3 | 142 | 1 |
| 4 | 27 | 1 | 4 | 243 | 1 | 4 | 756 | 2 |
| 5 | 16 | 0 | 5 | 303 | 1 | 5 | 241 | 1 |
| 6 | 397 | 0 | 6 | 238 | 0 | 6 | 775 | 0 |
| 7 | 698 | 0 | 7 | 179 | 0 | 7 | 1314 | 1 |
| 8 | 511 | 1 | 8 | 568 | 0 | 8 | 1269 | 1 |
| 9 | 607 | 0 | 9 | 523 | 2 | 9 | 900 | 0 |
| 10 | 690 | 1 | 10 | 451 | 0 | 10 | 1340 | 0 |
| 11 | 1057 | 0 | 11 | 712 | 0 | 11 | 1444 | 1 |
| 12 | 1064 | 0 | 12 | 571 | 3 | 12 | 1501 | 1 |
| 13 | 1075 | 0 | 13 | 620 | 0 | 13 | 1580 | 0 |
| 14 | 899 | 0 | 14 | 535 | 0 | 14 | 1550 | 0 |
| 15 | 1042 | 0 | 15 | 628 | 0 | 15 | 1633 | 1 |

**Table S11. Key parameter values used in the analysis of the *E. coli* adaptation experiment**. The condition-specific parameter values listed below –#Frames, #Smooth and Acc. Threshold– were automatically determined using a genetic algorithm and corresponding training sets (Tables S9 to S10). Values for *Tumbling Freq*. were calculated employing those condition-specific parameter values and the **Tumbling Frequencies** tab of TaxisPy. *Tumbling Freq*. refers to the culture's tumbling frequency during the first 10 seconds. Column *Nr. Traj.* contains the number of trajectories used in the calculations.

| Condition | Video ID | # Frames | # Smooth | Acc. Threshold | Tumbling Freq. | Nr. Traj. |
|---|---|---|---|---|---|---|
| Control @ 1 min | RPMSB_2 | 1 | 1 | 540 | 0.33 | 276 |
| Control @ 5 min | RPMSB_3 | 1 | 5 | 500 | 0.26 | 216 |
| Control @15 min | RPMSB_4 | 1 | 5 | 600 | 0.29 | 238 |
| Control @ 20 min | RPMSB_5 | 1 | 3 | 887 | 0.3 | 354 |
| Stimulated @ 1 min | RPMSB_26 | 2 | 5 | 320 | 0.01 | 420 |
| Stimulated @ 5 min | RPMSB_27 | 3 | 2 | 400 | 0 | 315 |
| Stimulated @ 10 min | RPMSB_42 | 3 | 5 | 600 | 0 | 137 |
| Stimulated @ 15 min | RPMSB_43 | 2 | 4 | 200 | 0.11 | 253 |
| Stimulated @ 20 min | RPMSB_30 | 1 | 2 | 600 | 0.25 | 86 |
| Stimulated @ 40 min | RPMSB_45 | 2 | 5 | 160 | 0.22 | 308 |

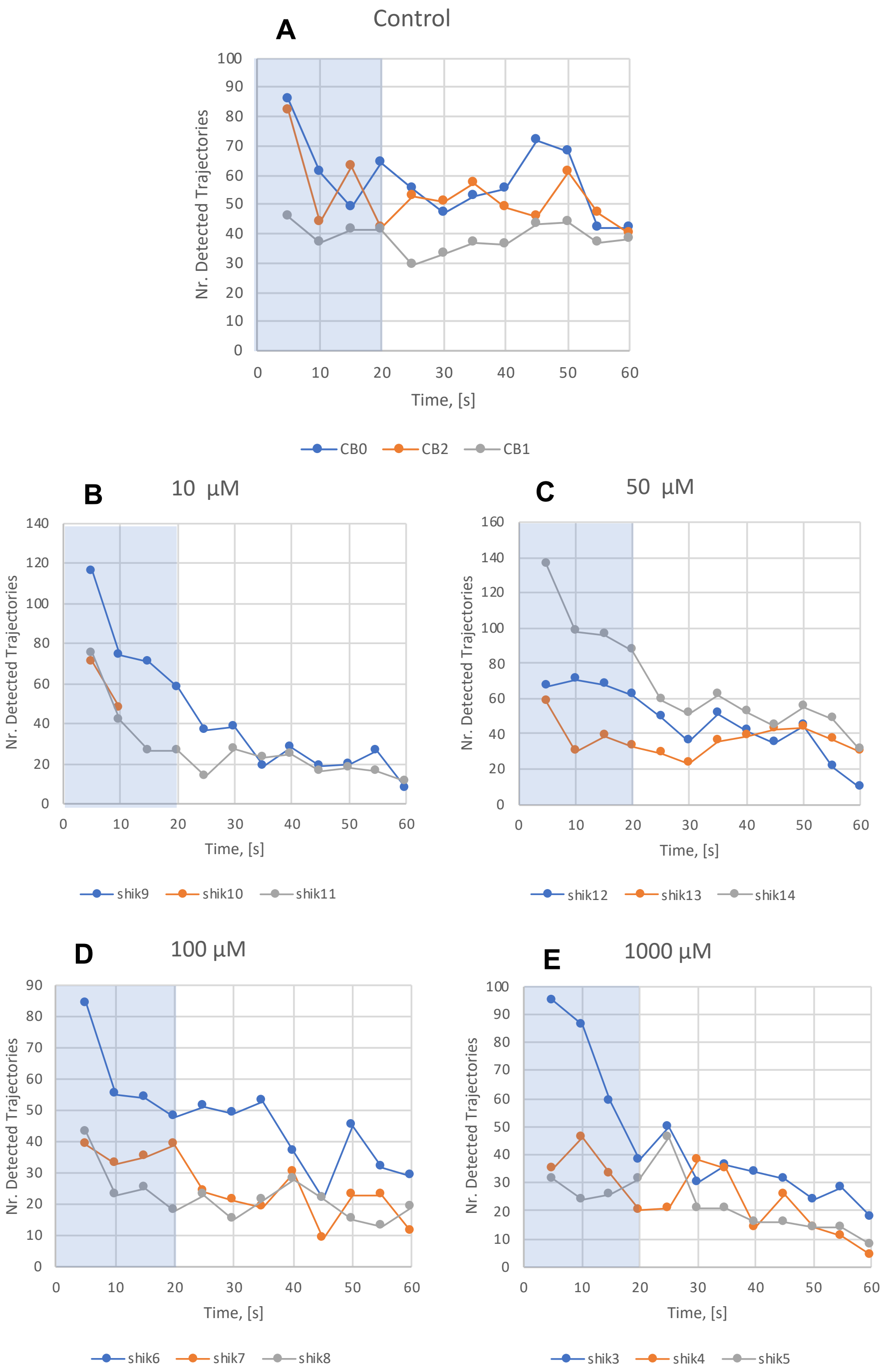


**Figure S1. Number of trajectories detected by TaxisPy as a function of time for the *P. putida* experiments**. Panel **A** shows the number of trajectories detected by TaxisPy as a function of time for the control condition. Panels **B** to **E** show the same temporal evolution for different concentrations of shikimate. The three lines in each panel correspond to each one of the triplicate videos recorded for each condition. The orange solid line in panel **B** is drawn only over ten seconds due to technical problems with the corresponding video which limited its length. The shaded blue areas represent the first 20 seconds of each video used to determine the minimum average tumbling frequency.
